# An in vivo drug screen reveals that cyclooxygenase 2-derived prostaglandin D_2_ promotes spinal cord neurogenesis

**DOI:** 10.1101/2023.10.05.561099

**Authors:** González-Llera Laura, Sobrido-Cameán Daniel, Quelle-Regaldie Ana, Sánchez Laura, Barreiro-Iglesias Antón

## Abstract

The study of neurogenesis is essential to understand fundamental developmental processes and for the development of cell replacement therapies for central nervous system disorders. Here, we designed an in vivo drug screening protocol in developing zebrafish to find new molecules and signalling pathways regulating neurogenesis in the ventral spinal cord. This unbiased drug screen revealed that 4 cyclooxygenase (COX) inhibitors reduce the generation of serotonergic interneurons in the developing spinal cord. These results fitted very nicely with available single cell RNAseq data revealing that the floor plate cells show differential expression of 1 of the 2 COX2 zebrafish genes (*ptgs2a)*. Indeed, several selective COX2 inhibitors and 2 different morpholinos against *ptgs2a* also caused a significant reduction in the number of serotonergic neurons in the ventral spinal cord and led to locomotor deficits. Single cell RNAseq data and different pharmacological manipulations further revealed that COX2-floor plate-derived prostaglandin D_2_ promotes neurogenesis in the developing spinal cord by promoting mitotic activity in progenitor cells. Rescue experiments using a phosphodiesterase-4 inhibitor suggest that intracellular changes in cAMP levels underlie the effects of COX inhibitors on neurogenesis and locomotion. Our study provides compelling in vivo evidence showing that prostaglandin signalling promotes neurogenesis in the ventral spinal cord.

## 1. Introduction

Studying neurogenesis, the process by which new neurons are produced from progenitor cells, is essential to understand central nervous system (CNS) development and for the implementation of cell replacement therapies for CNS disorders. Research on the spinal cord has been particularly valuable in uncovering the molecular mechanisms that control neurogenesis and the determination of cell fate from progenitor cells (for a review see Sagner and Briscoe, 2019).

Research on the spinal cord of various vertebrate species in recent decades has demonstrated that signalling molecules influence progenitor cells to develop distinct identities along the dorsoventral axis of the neural tube. For example, sonic hedgehog (Shh) signalling coming from the floor plate (FP) induces ventral identities in the spinal cord (Marti et al., 1995; Roelink et al., 1995). Shh induces the expression of specific transcription factors in progenitors of the ventral domains [from dorsal to ventral: the p0-2, pMN and p3 or lateral FP (LFP) domains]. Each of these progenitor domains gives rise to distinct neuronal subtypes, which are also characterized by the expression of specific transcription factors (see Sagner and Briscoe, 2019). For example, in zebrafish, serotonergic interneurons, which are generated from the LFP in the ventral spinal cord (Chen et al., 2023), are characterized by the expression of the transcription factor *pet1* (*fev*; Montgomery et al., 2016). Shh, and other morphogens, also play a role in regulating neurogenesis. For instance, in the pMN domain, high levels of Shh signalling initially drive symmetrical cell division, while decreasing Shh levels lead to differentiation of progenitor cells (Saade et al., 2013). Apart from the analysis of highly studied signalling pathways, like the Shh pathway, it would be of interest to find new molecules/signalling pathways regulating neurogenesis in the spinal cord. Deeper examination of the signalling pathways responsible for generating the diversity of neuronal subtypes from progenitor cells could uncover new targets for developing innovative treatments for movement disorders or spinal cord (or brain) injuries.

The zebrafish serves as a valuable model for uncovering new molecules and signalling pathways involved in spinal cord neurogenesis. This can be achieved thanks to the implementation of unbiased small molecule drug screens and thanks to available single cell RNAseq (scRNAseq) data from developing animals (e.g., Farnsworth et al., 2020). Zebrafish provide an ideal model for drug screens due to their rapid development, transparency of early embryos and larvae. Additionally, drugs can be applied in the water from where they are easily taken through the zebrafish skin (see Patton et al., 2021). Here, we developed a 2-step screening protocol to find new small molecules regulating spinal cord neurogenesis in developing zebrafish. For this, we decided to use as a model the population of serotonergic interneurons of the ventral spinal cord. This neuronal population offered a good model system to perform a drug screen due to the late appearance of these neurons during early development. Serotonergic neurons start to differentiate from 68 hours post-fertilization (hpf) (Montgomery et al., 2016), whereas, for example, most motor neurons are generated between 14 and 48 hpf (Barreiro-Iglesias et al., 2015). This allowed us to apply drugs at 48 hpf after the earlier developmental period, which supports the specificity of the drug screen results (the drugs will not affect early gross embryo morphogenesis).

Our drug screen revealed that several cyclooxygenase (COX) inhibitors decreased the generation of serotonergic interneurons in the developing zebrafish spinal cord. COXs (COX1 or COX2) are the rate-limiting enzymes in the generation of prostanoids [e.g., prostaglandins (PGs)]. COXs convert arachidonic acid to PGH_2_, which is then metabolized to other PGs, like PGE_2_ or PGD_2_, by specific PG synthases (see Nango and Kosuge, 2022. In the CNS, PGs are mainly known for their roles in pain, fever, apoptosis, or inflammatory processes (see Phillis et al., 2006), but there is very limited knowledge on their possible role in neurogenic processes in vivo, especially during early developing periods (see Barreiro-Iglesias, 2021). Our drug screen results fitted very nicely with available scRNAseq data showing that FP cells express 1 of the 2 COX2 zebrafish genes (*ptgs2a*) (Farnsworth et al., 2020). Indeed, specific COX2 inhibitors and *ptgs2a* morpholinos also reduced the number of serotonergic neurons in the spinal cord and led to locomotor deficits. ScRNAseq data also showed that FP spinal cord cells express 2 PGD_2_ synthase (PGDS) genes (*ptgdsb.1* and *ptgdsb.2*). Treatments with PGDS inhibitors also reduced the number of serotonergic interneurons in the spinal cord. Moreover, PGD_2_ rescued the effect of a COX inhibitor. Analyses of cell death and mitotic activity revealed that the reduction in serotonergic neurons after inhibiting PG synthesis is caused by decreased mitotic activity in progenitor cells and not by increased cell death. Rescue experiments using a phosphodiesterase-4 (PDE4) inhibitor suggest that intracellular changes in cAMP levels underlie the effects of COX inhibitors on neurogenesis and locomotor deficits. Overall, our results indicate that COX2-FP-derived PGD_2_ promotes neurogenesis in the ventral spinal cord by promoting the proliferation of progenitor cells. Our study is the first to provide compelling in vivo evidence for a role of PGs in the regulation of the neurogenic process in developing animals. Furthermore, our study provides a new drug screen protocol to find small molecules regulating spinal cord neurogenesis in developing vertebrates.

## 2. Material and methods

### 2.1 Animals

All zebrafish lines were kept and raised under standard conditions (Westerfield, 2000) in the fish facilities of the Department of Genetics of the University of Santiago de Compostela (code of the facility: AE-LU-003, ES270280346401). All experiments were approved by the Bioethics committee of the University of Santiago de Compostela and the Xunta de Galicia (project license no.: 01/20/LU-003) and were carried out in accordance with EU Directive 2010/63/EU for animal experiments. For experimental analyses, we used wild type or transgenic Tg(−3.2*fev*:EGFP) [referred also as *pet1*:gfp; obtained from the European Zebrafish Resource Center (EZRC), EZRC code ne0214Tg] larvae [up to 4 days post-fertilization (dpf)]. Embryos were collected from the breeding tanks and were divided into Petri dishes at a density of maximally 100 embryos per dish until they were 2 dpf (when they were used for drug treatments, see below), but no formal randomization method was used. For this study a total of 3460 zebrafish embryos were used. The specific number of animals used for each experiment is indicated in the figure legends or in the supplementary files.

### 2.2 Drug screen

We carried out an unbiased drug screen of the LOPAC®1280 – Small Scale library (Sigma; Cat#LO4200-1EA; International Version). We screened drugs from racks 9 and 10 of the library (160 drugs in total). In the library, drugs are diluted in DMSO at stock concentration of 10 mM. For treatments, 2 µl of the stock solution were diluted in 2 ml of fish water (reverse osmosis-purified water; working dilution of 10 µM). Larvae were incubated from 2 dpf to 4 dpf in groups of 5 animals per well in 24 well-plates (Supplementary Figure 1). Control animals were always treated with DMSO alone.

We also tested selective COX2 inhibitors present in other racks of the LOPAC® library: Etodolac, Rofecoxib, Nimesulide, Niflumic Acid and DFU. These treatments were carried out as with the other drugs of the library in the unbiased drug screen.

### 2.3 Treatments with specific drugs in whole larvae

Specific drugs (see Table 1) were diluted in DMSO at stock concentration of 10 mM. For treatments, 2 µl of the stock solution were diluted in 2 ml of fish water (working dilution of 10 µM). Larvae were incubated from 2 dpf until 3 or 4 dpf in groups of 5 animals per well in 24 well-plates (Supplementary Figure 1A). Control animals were always treated with DMSO alone.

**Table 1.**
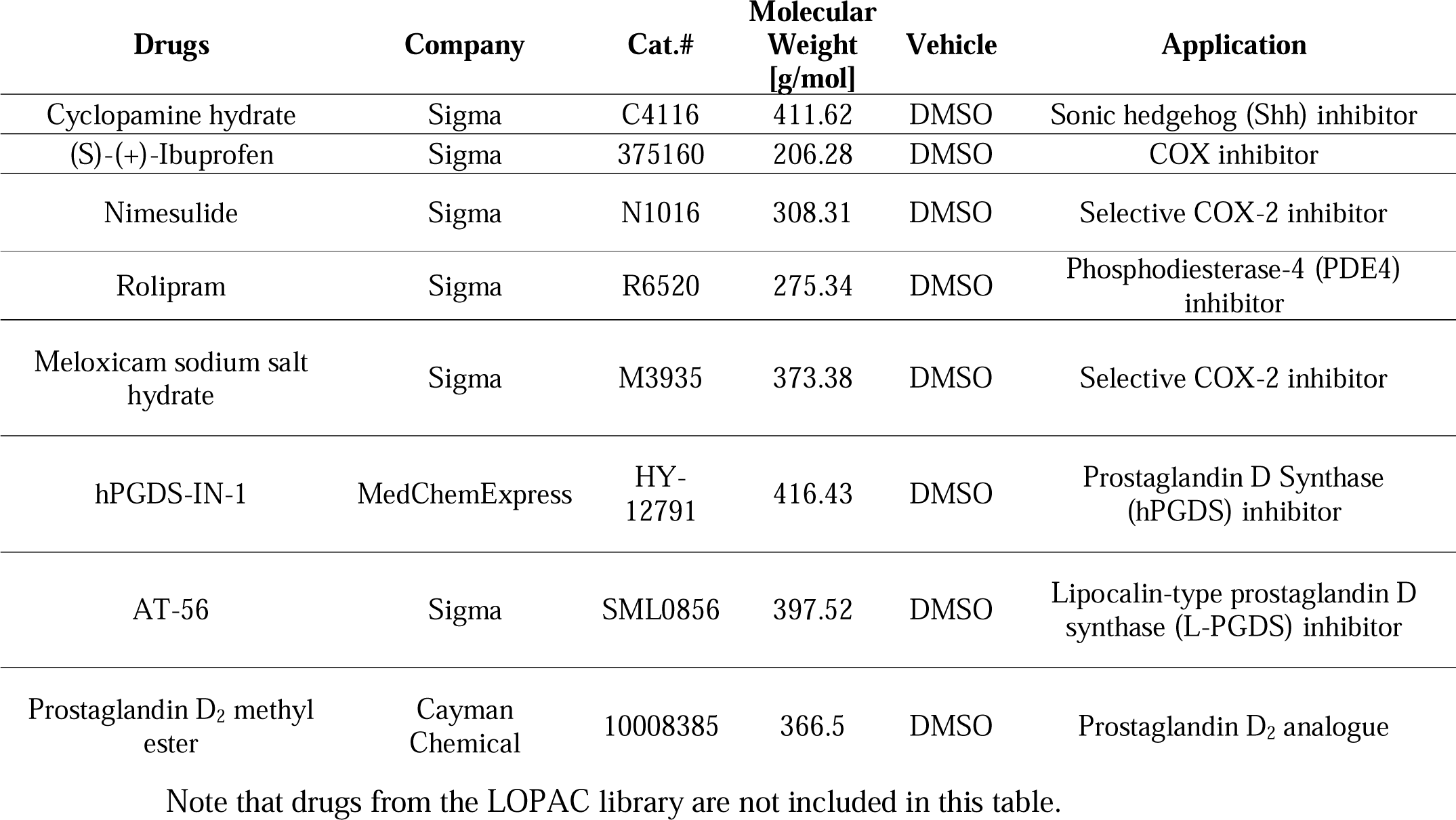
Drugs used in specific experiments.

### 2.4 Behavioural analyses

Locomotor performance of 4 dpf zebrafish larvae after drug treatments was quantified with the Zebralab software using a Zebrabox (Viewpoint; Civrieux, France). The quantification software measures the distance moved by each fish in a certain period. For this analyses, 4 dpf larvae were transferred to 96 well-plates (1 animal per well). Before measuring locomotor activity, the larvae were left in clean fish water without drugs or DMSO (control group) for 1 hour. Total larval movement was measured for 1 hour, alternating 10-minutes periods of light and dark conditions. Larvae were kept at a constant standard temperature of 28.5° C while measuring locomotor activity. The locomotor activity of each larva was calculated based on the distance moved (in cm) in the 6 10-minute periods.

### 2.5 Morpholino treatments

Morpholinos (Gene Tools, LLC) diluted in nuclease-free water were injected (approximately 3 nl; 1 mM) into the yolk at the one-cell stage of development. Control animals were injected with the Standard Control morpholino (5’- CCTCTTACCTCAGTTACAATTTATA-3’) from Gene Tools. To knockdown the expression of *ptgs2a,* animals were injected with previously designed *ptgs2a* translation blocking (5’- AACCAGTTTATTCATTCCAGAAGTG-3’; Grosser et al., 2002; ZFIN ID: ZDB-MRPHLNO-050722-5; ZFIN name: MO1-ptgs2a) or splicing (5’- ATTCAACTTACACAACAGGATATAG - 3’; Yeh et al., 2009; ZFIN ID: ZDB-MRPHLNO-110427-4; ZFIN name: MO5-ptgs2a) morpholinos. After morpholino administration, larvae were left until 4 dpf in fish water.

### 2.6 Whole-mount anti-serotonin (5-HT) or anti-gfp immunofluorescence

After drug or morpholino treatments, 4 dpf larvae were euthanized by tricaine methanesulfonate (Sigma) overdose and then fixed with 4% paraformaldehyde (PFA) in phosphate-buffered saline (PBS; pH 7.4) for 2 h at 4°C. After washes in PBS, 4 dpf larvae were incubated in Proteinase K from *Tritirachium album* (Sigma; Cat#: P4850; ≥800U/mL; 1 µL per ml of PBS) for 35 min at room temperature and then in glycine [50 mM in PBS with 0,2% Triton X-100 (PBST)] for 10 min at room temperature. Then, the larvae were incubated with rabbit anti-5-HT (Immunostar, Still Water, MN, USA; Cat#: 20080; dilution 1:2500; RRID:AB_572263) or chicken anti-gfp (Abcam, Cambridge, UK; Cat#: ab13970; dilution 1:500; RRID:AB_300798) antibodies overnight at 4°C. Then, they were rinsed in PBST and incubated overnight at 4°C with Cy3-conjugated goat anti-rabbit (Jackson ImmunoResearch; Cat#: 111-165-144; dilution 1:500; RRID:AB_2338006) or Alexa Fluor 488-conjugated goat anti-chicken (Thermo Fisher Scientific; Waltham, MA, USA; Cat#: A-11039; dilution 1:500; RRID:AB_2534096) antibodies. Antibodies were always diluted in PBST with 1% DMSO, 1% normal goat serum and 1% bovine serum albumin. Larvae were mounted with 70% glycerol in PBS. Control and treated animals were always processed in parallel and the same antibody solutions were used for both control and treated animals in each experiment.

### 2.7 Cryostat sections

3 dpf larvae were euthanized by tricaine methanesulfonate (Sigma) overdose and then fixed with PFA in PBS (pH 7.4) for 2 h at 4°C. After rinsing in PBS, larvae were cryoprotected overnight with 30% sucrose in PBS, embedded in Neg-50^TM^ (Thermo Scientific, Kalamazoo, MI, USA), and frozen with liquid nitrogen-cooled isopentane. Transverse sections of the body starting from the caudal fin (16 μm thickness) were obtained on a cryostat and mounted on Superfrost Plus slides (Menzel-Glasser, Madison, WI, USA).

### 2.8 TUNEL labelling on cryostat sections

TUNEL staining was performed according to the manufacturer’s protocol with minor modifications (In Situ Cell Death Detection Kit, TMR red; catalogue number 12156792910; Roche, Mannheim, Germany). Briefly, the sections were incubated in methanol for 15 min at −20°C to permeabilize the lipid membranes, followed by brief washes in PBS and another incubation in 0.01 M citrate buffer pH 6.0 for 30 min at 70°C. After several washes in PBS, sections were incubated in the TUNEL reaction mix, containing the Labeling Solution (TMR red labeled nucleotides) and the Enzyme Solution (terminal deoxynucleotidyl transferase), for 90 min at 37°C. Slides were washed in PBS and distilled water, allowed to dry for 30 min at 37°C, and mounted with MOWIOL® 4-88 (Calbiochem, Darmstadt, Germany). Negative controls were obtained by incubating sections only with the Labeling Solution (without terminal deoxynucleotidyl transferase). Positive controls were generated by incubating some sections from control 3 dpf animals with recombinant DNAse I (400 U/ml; Roche) for 20 minutes at room temperature before the incubation in the reaction mix. Sections from control and drug treated animals were always processed in parallel and the same TUNEL labeling solution was used for sections from control and drug treated animals.

### 2.9 Immunofluorescence on cryostat sections

Sections were first treated with 0.01 M citrate buffer pH 6.0 for 30 min at 90°C for heat-induced epitope retrieval, allowed to cool for 10 min in cold water, and then rinsed in 0.05 M Tris-buffered saline (TBS) pH 7.4 for 20 min. Then, the sections were incubated overnight at RT with a rabbit polyclonal anti-pH3 antibody (1:500; Sigma; Cat#: H0412; RRID: AB_477043). Sections were then rinsed 3 times in TBS for 15 min each and incubated for 1 hour at room temperature with a Cy3-conjugated goat anti-rabbit IgG antibody (Jackson ImmunoResearch; Cat#: 111-165-144; dilution 1:500; RRID:AB_2338006). All antibody dilutions were made in TBS containing 15% normal goat serum (Millipore) and 0.2% Triton X-100 (Sigma). Finally, sections were rinsed 3 times in TBS for 15 min each and in distilled water for 20 min, allowed to dry for 30 min at 37 °C, and mounted in MOWIOL^®^ 4-88 (Calbiochem). Sections from control and treated animals were always processed in parallel for each antibody staining and the same antibody solutions were used for sections of control and treated animals in each experiment.

### 2.10 Imaging and cell counting in whole-mounted larvae and spinal cord sections

After anti-5-HT immunofluorescence experiments in whole-mounted 4 dpf larvae, confocal photomicrographs were taken at the level of the caudal fin with TCS-SP2 spectral or Stellaris 8 confocal laser microscopes (Leica Microsystems) with a 20x objective. For the quantification of serotonergic neurons, the total number of 5-HT-immunoreactive (ir) interneurons located at the level of the caudal fin was quantified manually going through the stack of confocal optical sections. The experimenter was blinded during quantifications.

After immunofluorescence experiments or TUNEL labeling in cryostat transverse sections, confocal photomicrographs were taken with the Stellaris 8 confocal laser microscope (Leica Microsystems). We quantified the number of labelled cells in 1 out of each 3 consecutive spinal cord transverse sections starting from the caudal end of the spinal cord and moving rostrally. 9 sections were quantified in each animal and then the mean number of labelled cells per section was calculated for each animal. For pH3 immunolabeling we quantified separately the number of positive cells in the ventral and dorsal portions of the spinal cord.

After cell quantifications, figures were prepared with Adobe Photoshop 2022 (San Jose, CA, USA) with minor adjustments of brightness and contrast of the confocal images. Schematic drawings were also generated with Adobe Photoshop 2022.

### 2.11 Statistical analysis

Experiments with locomotor analyses or cell quantifications were carried out in a minimum of 3 different clutches of animals for each different drug or morpholino treatment, and a minimum of 15 animals were included in each experimental group. Each dot in the graphs represents one animal and n numbers for each experimental group are indicated in the figure legends or in the supplementary files. Statistical analyses were performed with Prism 9 (GraphPad software, La Jolla, CA, USA). Normality of the data was determined with the D’Agostino & Pearson test. For groups with low n number in the drug screen we used a Shapiro-Wilk normality test. To determine statistically significant differences (p ≤ 0.05) between two groups of normally distributed data we used an unpaired (Student’s) t-test (two-tailed). To determine statistically significant differences between two groups of non-normally distributed data we used a Mann Whitney U test (two-tailed). To determine statistically significant differences between three groups of non-normally distributed data we used a Kruskal-Wallis test and *post hoc* Dunn’s multiple comparison test. In the figures, significance values were represented by a different number of asterisks in the graphs: *p-value between 0.01 and 0.05, **p-value between 0.001 and 0.01, ***p-value between 0.001 and 0.0001, ****p-value <0.0001. Exact p-values are given in the figure legends.

## 3. Results and discussion

### 3.1 An unbiased drug screen reveals that several COX inhibitors reduce the number of serotonergic cells in the spinal cord

Spinal cord serotonergic cells begin to differentiate in developing zebrafish at around 60 hpf, when *pet1 (fev)* expression is first observed. The first mature cells expressing *tryptophan hydroxylase 2* (the rate-limiting enzyme for 5-HT synthesis) can be observed at 68 hpf and a fully mature 5-HT-ir population can be observed at 4 dpf (Montgomery et al., 2016). Based on this developing/differentiating timing we decided to apply drugs in our drug screen at 2 dpf, and for 2 days (Supplementary Figure 1A, B), aiming to identify drugs that could affect the behaviour of progenitor cells and/or the neuronal differentiation process. As indicated above, applying the drugs at this time point allowed us to avoid the earlier gross developmental period and other major ventral neurogenic events like the earlier period of motor neuron generation.

We designed a two-step drug screening protocol (see Supplementary Figure 1B). In the primary screen, each drug from the LOPAC^®^ library was tested (at 10 µM) in 10 larvae (2 wells with 5 animals per well). In the secondary screen, hits from the primary screen (drugs that significantly changed the number of serotonergic interneurons as compared to DMSO controls) were further tested in groups of 15 to 30 larvae (Supplementary Figure 1B).

First, we used Cyclopamine (a Shh signalling inhibitor) as a positive control to confirm the validity of our drug screening protocol. There was no previous data on the role of Shh signalling in the generation of spinal serotonergic neurons in developing zebrafish. However, the ventral location and origin from the LFP of these neurons (Montgomery et al., 2016, 2018; Chen et al., 2023) suggests that their generation is probably influenced by Shh signalling. Moreover, after spinal cord injury in adult zebrafish, these serotonergic neurons are also regenerated from the p3/LFP domain and a Cyclopamine treatment inhibits their regeneration (Kuscha et al., 2012). Indeed, following this drug treatment protocol we observed a significant reduction in the number of 5-HT-ir neurons in Cyclopamine treated larvae as compared to DMSO controls (Fig. 1A). This indicates that Shh signalling regulates the generation of serotonergic neurons in developing animals as during neuronal regeneration in adult zebrafish (Kuscha et al., 2012), and confirms the validity of our screening protocol to find new small molecules affecting ventral neurogenesis in the developing spinal cord.

**Figure 1.**
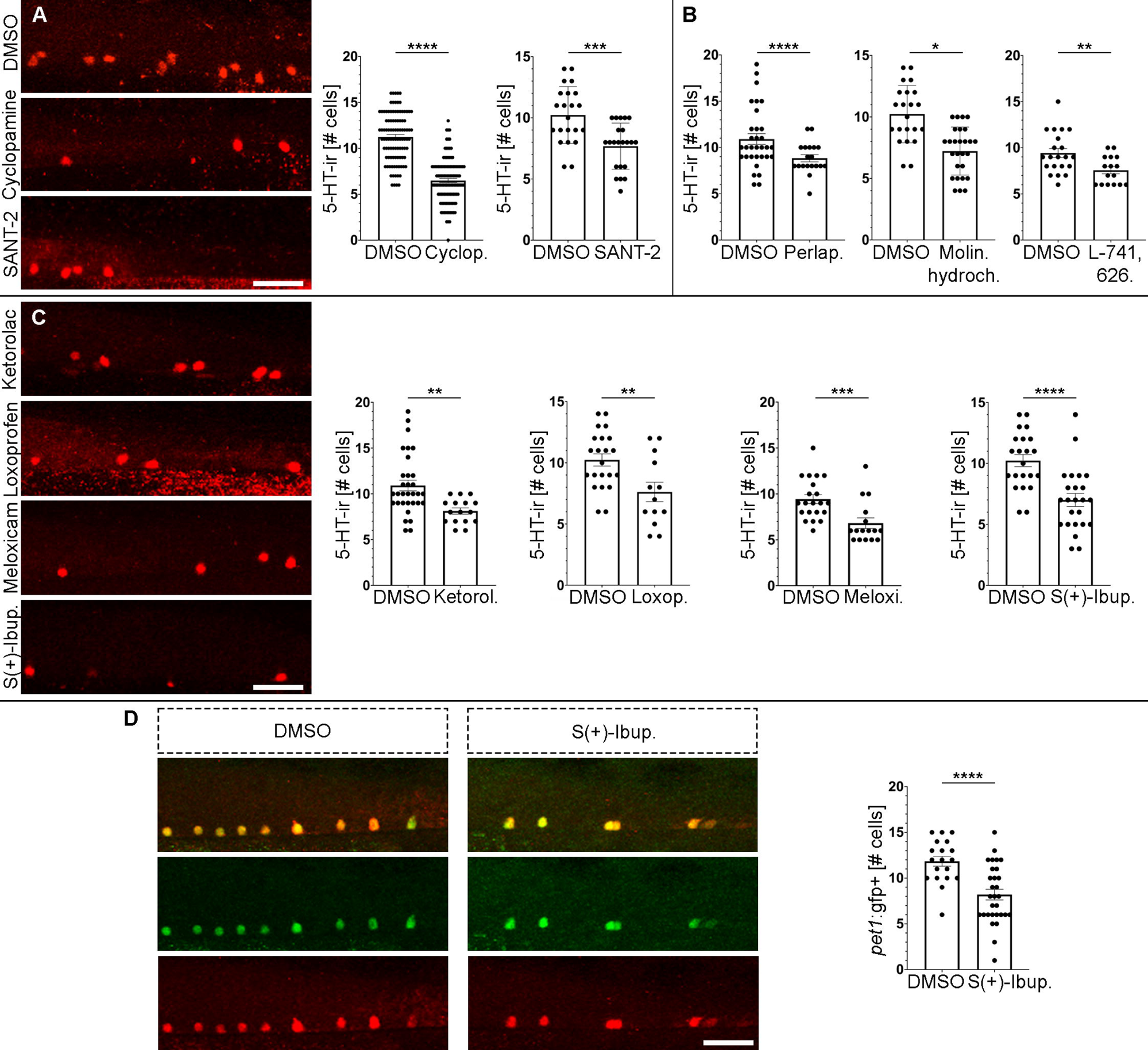
An unbiased drug screen reveals new small molecules and signalling pathways that control neurogenesis in the ventral spinal cord. A. Treatments with the Shh signalling inhibitor Cyclopamine (6.487 ± 0,224 cells, n=117; Mann-Whitney test; p-value <0.0001) or the smoothened receptor antagonist SANT-2 (7.682 ± 0.402 cells, n=22; Unpaired t test; p-value = 0.0003) reduced the numbers of serotonergic neurons (red fluorescence) in the ventral spinal cord as compared to DMSO controls (Cyclopamine control: 11.24 ± 0.264 cells, n=101; SANT-2 control: 10.23 ± 0.496 cells, n=22). B. Treatments with anti-dopaminergic drugs from the LOPAC library (Perlapine: 7.222 ± 0.375 cells, n=27; Unpaired t test; p-value <0.0001; Molindone Hydrochloride: 8.842 ± 0.384 cells, n=19; Unpaired t test; p-value = 0.0142; L-741,626: 7.563 ± 0.376 cells, n=21; Unpaired t test; p-value = 0.0067) reduced the numbers of serotonergic neurons in the ventral spinal cord as compared to DMSO controls (Perlapine control: 10.23 ± 0.496 cells, n=22; Molindone Hydrochloride control: 10.91 ± 0.581 cells, n=32; L-741,626 control: 9.429 ± 0.486 cells, n=16). C. Treatments with COX inhibitors from the LOPAC library (Ketorolac: 8.125 ± 0.340 cells, n=16; Unpaired t test; p-value = 0.0022; Loxoprofen: 7.615 ± 0.789 cells, n=13; Unpaired t test; p-value = 0.0058; Meloxicam: 6.813 ± 0.579 cells, n=16; Mann-Whitney test; p-value = 0.0005; S(+)-Ibuprofen: 7.000 ± 0.532 cells, n=24; Unpaired t test; p-value <0.0001) reduced the numbers of serotonergic neurons in the ventral spinal cord as compared to DMSO controls (Ketorolac control: 10.91 ± 0.581 cells, n=32; Loxoprofen control: 10.23 ± 0.496 cells, n=13; Meloxicam control: 9.429 ± 0.486 cells, n=21; S(+)- Ibuprofen control: 10.23 ± 0.496 cells, n=22). D. A treatment with the COX inhibitor S(+)-Ibuprofen (8.200 ± 0.582 cells, n=30; Unpaired t test; p-value <0.0001) reduced the numbers of serotonergic pet1+ neurons in the ventral spinal cord of *pet1*:gfp fish as compared to DMSO controls (11.84 ± 0.542 cells, n=19). Note the perfect colocalization of serotonin (red) and gfp (green) immunofluorescence signals in serotonergic cells. Rostral is to the right and dorsal to the top in all photomicrographs. Scale bars: 25 µm.

We screened 160 compounds of the LOPAC^®^ library (racks 9 and 10) in the primary screen (Supplementary File 1). Cyclopamine was included as a positive control in all the drug screen experiments and always caused a significant reduction in numbers of serotonergic neurons (Supplementary File 1). 7 drugs killed more than 30% of the animals and were discarded for the secondary screen (Supplementary File 1). Of the remaining 153 compounds, 40 drugs significantly reduced the number of 5-HT-ir neurons at 4 dpf as compared to DMSO controls (Supplementary File 1). In the secondary screen, with larger groups of animals, we confirmed that 26 of these 40 compounds significantly reduce the number of 5-HT-ir neurons (Supplementary File 2). We found one discrepancy between the primary and secondary screens with the drug TMPH Hydrochloride (an antagonist of nicotinic acetylcholine receptors), which caused a significant increase in 5-HT-ir neurons in the secondary screen (Supplementary File 2). Interestingly, among the 26 drugs reducing the number of 5-HT-ir neurons in the secondary screen there was another Shh signalling inhibitor (SANT-2; a smoothened receptor antagonist; Fig. 1A), which confirms that Shh signalling regulates the generation of serotonergic spinal cord interneurons. In addition, 3 of the 26 drugs reducing the number of 5-HT-ir neurons have anti-dopaminergic activity (Perlapine, Molindone Hydrochloride and L-741,626; Fig. 1B), which fits nicely with previous data showing that dopamine promotes motor neuron generation in the developing zebrafish spinal cord (Reimer et al., 2013). Overall, these results confirm the validity of our screening protocol and reveal that Shh and dopamine regulate the generation of serotonergic neurons as previously shown for spinal cord motor neurons.

Among the other 22 drugs reducing the number of 5-HT-ir neurons, there were 4 COX inhibitors (Ketorolac Tris Salt, Loxoprofen, Meloxicam Sodium and S(+)- Ibuprofen; Fig. 1C; Supplementary File 2). These unbiased drug screen results pointed to a previously unknown role of COXs and prostanoids in promoting neurogenesis in the ventral spinal cord. However, to confirm that COX inhibitors inhibit neurogenesis and not 5-HT production in existing cells, we repeated the S(+)-Ibuprofen treatments in a *pet1*:gfp transgenic zebrafish line. The S(+)-Ibuprofen (10 µM) treatment starting at 2 dpf significantly reduced the number of *pet1*:gfp+ cells in the spinal cord of 4 dpf zebrafish (Fig. 1D). Therefore, COX-derived prostanoids promote the generation of serotonergic neurons in the ventral spinal cord.

Interestingly, the unbiased drug screen results also suggested that the reduction in serotonergic neurons could be related to COX2 inhibition (as opposed to COX1 inhibition). S(+)-Ibuprofen (the active enantiomer of ibuprofen), Ketorolac and Loxoprofen are non-selective COX inhibitors, but Meloxicam Sodium shows 300-fold selectivity for COX2. In addition, among the non-significant drugs in the primary screen, there were 2 selective COX1 inhibitors: Ketoprofen (p = 0.0977) and Indomethacin (p = 0.5940) (Supplementary File 1). Overall, our drug screen results pointed to a possible role for COX2 in promoting neurogenesis in the ventral spinal cord.

### 3.2 Selective COX2 inhibitors reduce the number of serotonergic cells in the spinal cord

Based on the unbiased drug screen results suggesting a role for COX2 in promoting spinal cord neurogenesis, we decided to analyse the effect of 5 selective COX2 inhibitors (Etodolac, Rofecoxib, Nimesulide, Niflumic Acid and DFU) available in other racks of the LOPAC^®^ library and that were not included in the unbiased drug screen (Supplementary File 3). Treatments with Etodolac, Rofecoxib, Nimesulide and DFU (all at 10 µM) starting at 2 dpf significantly reduced the number of 5-HT-ir neurons in the spinal cord of 4 dpf zebrafish (Fig. 2A; Supplementary File 3). Niflumic Acid also reduced the number of 5-HT-ir neurons at 4 dpf, but it caused a high mortality (Supplementary File 3), which could be related to other effects of the drug as a chloride or calcium channel inhibitor (Balderas et al., 2012). These results with selective COX2 inhibitors confirmed that COX2 activity promotes neurogenesis in the ventral spinal cord.

**Figure 2.**
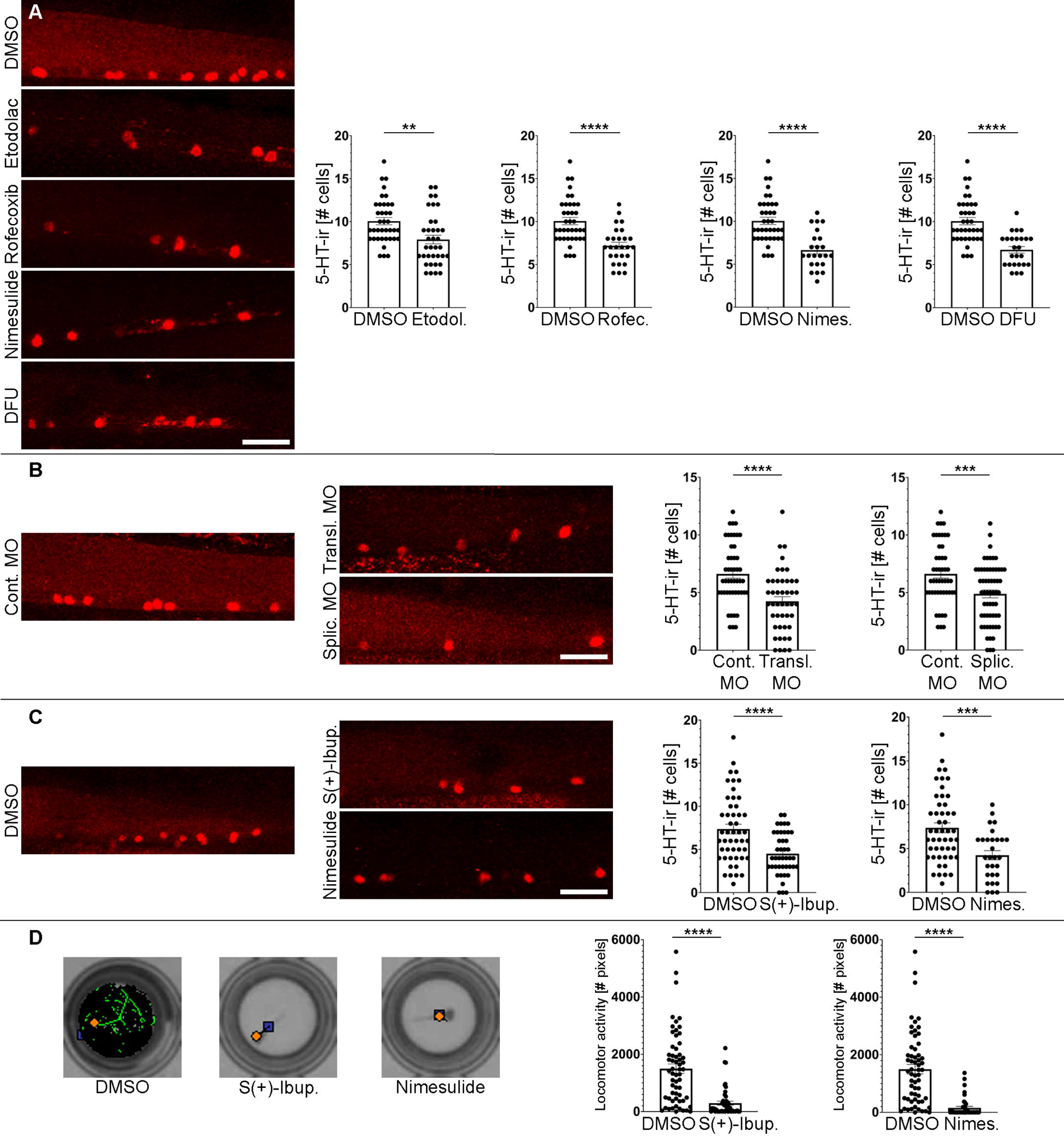
COX2 (*ptgs2a*) inhibition reduces the numbers of serotonergic neurons (red fluorescence) in the ventral spinal cord and causes locomotor deficits. A. Treatments with the selective COX2 inhibitors Etodolac (7.917 ± 0.499 cells, n=36; Unpaired t test; p-value = 0.0017), Rofecoxib (7.185 ± 0.403 cells, n=27; Unpaired t test; p-value <0.0001), Nimesulide (6.652 ± 0.46 cells, n=23; Unpaired t test; p-value <0.0001) and DFU (6.72 ± 0.38 cells, n=25; Unpaired t test; p-value <0.0001) reduced the numbers of serotonergic neurons in the ventral spinal cord as compared to DMSO controls (10.05 ± 0.424 cells, n=38). B. Translation (4.244 ± 0.395 cells, n=45; Unpaired t test; p-value = 0.0007) and splicing (4.881 ± 0.338 cells, n=59; Unpaired t test; p-value = 0.0015) morpholinos (MO) against the *ptgs2a* mRNA reduced the numbers of serotonergic neurons in the ventral spinal cord as compared to zebrafish treated with the control morpholino (6.250 ± 0.411 cells, n=40). C. Treatments with the COX inhibitors S(+)- Ibuprofen (4.512 ± 0.375 cells, n=43; Unpaired t test; p-value <0.0001) and Nimesulide (4.241 ± 0.519 cells, n=29; Unpaired t test; p-value = 0.0003) reduced the numbers of serotonergic neurons in the ventral spinal cord as compared to DMSO controls (7.367 ± 0.563 cells, n=49). D. Animals treated with S(+)-Ibuprofen (294.2 ± 69.20 cm, n=49; Mann-Whitney test; p-value <0.0001) and Nimesulide (155.5 ± 50.63 cm, n=42; Mann-Whitney test; p-value <0.0001) (see C) showed significant locomotor deficits as compared to DMSO controls (1,499 ± 163.1 cm, n=60). Examples of 10-minute swim tracks (with light) are shown to the left. Total locomotor activity, which was recorded for 1 hour (6 10-minute periods alternating light and dark conditions), is shown in the graphs. Rostral is to the right and dorsal to the top in all photomicrographs. Scale bars: 25 µm.

### 3.3 ScRNAseq data from developing zebrafish reveals differential expression of a COX2 gene (ptgs2a) in FP cells

An obvious cell population as a possible source of prostanoids for the regulation of neurogenesis in the ventral spinal cord is the FP, a specialized glial structure of the ventral midline mainly known for its role in ventral nervous tissue differentiation through Shh secretion (see introduction). To determine if zebrafish FP cells express any of the zebrafish COX genes (*ptgs1*, *ptgs2a* or *ptgs2b*), we used an available single cell transcriptome whole-body atlas from developing zebrafish (Farnsworth et al., 2020). This atlas was generated from 1, 2 and 5 dpf zebrafish, which nicely covers the developmental period for the generation of serotonergic neurons. In the atlas, cell cluster 176 was identified as the population of spinal cord FP cells based on the differential expression of several well-known FP marker genes (e.g., *shha*, *shhb*, *gfap*, *slit1a*, *slit2*, *slit1b*, *foxj1a*, *wnt4b*, *spon1a, spon1b* or *ctgfa*) (Farnsworth et al., 2020). Importantly, one of the genes showing differential expression in this cluster was *ptgs2a* (p-value = 7.38E-56; adjusted p-value = 2.4E-51). 51 out of the 94 cells (54.25%) assigned to the spinal cord FP cluster (cluster number 176 of the atlas) show *ptgs2a* expression (Supplementary Figure 2A). Therefore, *ptgs2a* is differentially expressed in a cell population that is in an optimal location to influence the generation of serotonergic cells in the developing spinal cord, as with Shh signalling coming also from the floor plate (see above).

Based on the single cell transcriptomic data, we decided to manipulate *ptgs2a* expression by using well-characterized translation (Grosser et al., 2002) and splicing (Yeh et al., 2009) morpholinos against the zebrafish *ptgs2a* transcript. These morpholinos had been previously used to knockdown *ptgs2a* expression in developing zebrafish (e.g., Poureetezadi et al., 2016; Marra et al., 2019). For example, *ptgs2a* knockdown efficacy with the splicing morpholino in zebrafish was previously demonstrated by RT-PCR (Poureetezadi et al., 2016). Administration of translation blocking or splicing morpholinos significantly reduced the numbers of 5-HT-ir neurons in the ventral spinal cord at 4 dpf as compared to animals that received the control morpholino (Fig. 2B). This confirmed that the effects of the COX2 inhibitors are probably caused by *ptgs2a* inhibition and that *ptgs2a*-derived prostanoids promote neurogenesis in the ventral spinal cord. In previous work, only a few studies reported that COX2 inhibition affects adult neurogenesis in rodents (Goncalves et al., 2010; Nam et al., 2013, 2015; for a review see Barreiro-Iglesias, 2021). Meloxicam and Nimesulide treatments decreased the appearance of new neurons in the olfactory bulb of 6-week-old mice (Goncalves et al., 2010). A Celecoxib (a COX2 inhibitor) treatment also reduced numbers of doublecortin+ neuroblasts in the dentate gyrus of 9-week-old mice (Nam et al., 2015). COX2 knockout 8-week-old mice also exhibit a significant reduction in doublecortin+ neuroblasts of the dentate gyrus (Nam et al., 2013, 2015). Thus, our study is the first to reveal a role for COX2 (*ptgs2a*) in promoting neurogenesis in the spinal cord and during early developmental periods.

### 3.4 Neurogenesis inhibition by non-steroidal anti-inflammatory drugs (NSAIDs) leads to locomotor deficits

8 different COX inhibitors and *ptgs2a* knockdown with 2 different morpholinos caused a clear reduction in numbers of serotonergic spinal cord neurons of 4 dpf zebrafish (see above). Recent work has shown that serotonergic signalling from these intrinsic spinal cord neurons regulates locomotion by reducing spinally produced motor-bursting in zebrafish (Montgomery et al., 2018). Consequently, it is of interest to test whether the reduction in the generation of serotonergic neurons has an impact on the locomotor activity of the 4 dpf larva. We measured locomotor activity during 1 hour (6 10-minute periods alternating light and dark conditions) in control and S(+)-Ibuprofen or Nimesulide treated (2 to 4 dpf) zebrafish. 4 dpf zebrafish were left in 96-well plates (1 larva per well) without DMSO or NSAIDs for 1 hour before measuring locomotor performance with the Zebrabox system (see Material and Methods). First, we confirmed that the drugs (these were newly purchased and not from the LOPAC^®^ library) also caused a significant reduction in numbers of 5-HT-ir spinal cord neurons in 4 dpf zebrafish as compared to DMSO controls (Fig. 2C). Notably, these treatments led to a significant reduction in locomotor activity during the hour of swim tracking (Fig. 2D). Thus, the decrease in spinal cord neurogenesis due to COX2 inhibition leads to behavioural (locomotor) deficits in 4 dpf zebrafish. We should consider that this locomotor deficit could also be caused by changes in other mature cell types apart from serotonergic neurons. For example, during this developmental period pMN progenitors begin to generate oligodendrocytes after the initial developmental period dedicated to motor neuron production (Park et al., 2005; Czopka et al., 2013; Ravanelli and Appel, 2015). Interestingly, a recent study has shown that PGDS derived from oligodendrocyte progenitor cells promotes oligodendrocyte development in mice (Pan et al., 2023). Thus, future work should investigate whether oligodendrogenesis is also regulated by COX2 signalling in the ventral spinal cord, which could also contribute to the locomotor deficits observed after COX2 inhibition.

### 3.5 PGD_2_ promotes spinal cord neurogenesis

COX2 converts arachidonic acid to PGH_2_, which is then metabolized to other PGs by specific PG synthases. To identify the specific PG or PGs promoting neurogenesis in the ventral spinal cord, we looked for PG synthases showing differential gene expression in the spinal cord FP cluster of the single cell transcriptome atlas (Farnsworth et al., 2020; see above). Indeed, spinal cord FP cells show differential expression of *ptgdsb.2* (p = 6.05E-08; adjusted p value = 0.00196786), which is one of the 3 PGDS genes (*ptgdsa, ptgdsb.1* and *ptgdsb.2*) in zebrafish. *Ptgdsb.2* expression is present in 27 out of the 94 cells (28.72%) assigned to the spinal cord FP cluster (cluster number 176 of the atlas; Supplementary Figure 2B; Farnsworth et al., 2020). Although no other PG synthase showed differential gene expression, spinal cord FP cells also express the *ptgdsb.1* gene (20 out of 94 of FP cells in cluster 176; Supplementary Figure 2C). Thus, based on the single cell transcriptomic data, we decided to treat 2 dpf animals with the PGDS inhibitors hPGDS-IN-1 or AT-56. Treatments with both PGDS inhibitors caused a significant reduction in the numbers of 5-HT-ir spinal cord neurons of 4 dpf zebrafish as compared to DMSO controls (Fig. 3A). Then, aiming to perform a gain of function experiment, we treated 2 dpf animals with the PGD_2_ analogue PGD_2_ methyl ester. In pilot experiments we observed pericardial/yolk sac edemas in most animals treated with PGD_2_ methyl ester at 10 or 5 µM (not shown), therefore for subsequent experiments we went down to a 2.5 µM concentration. In 4 dpf zebrafish treated with 2.5 µM PGD_2_ methyl ester, we did not observe a significant increase in the numbers of 5-HT-ir spinal cord neurons as compared to DMSO controls (Fig. 3A). However, the same treatment with 2.5 µM PGD_2_ methyl ester was able to rescue the inhibitory effects of an S(+)-Ibuprofen treatment (at 10 µM) on the generation of 5-HT-ir spinal cord neurons (Fig. 3B). These results could suggest that the ventral spinal cord is working at its maximum capacity for the generation of serotonergic neurons, which would explain why the PGD_2_ methyl ester treatment rescues the effects of a COX inhibitor but does not promote an increase in neurogenesis on its own. Overall, transcriptomic, and pharmacological data indicate that PGDS/PGD_2_ signalling from FP cells promotes neurogenesis in the ventral spinal cord.

**Figure 3.**
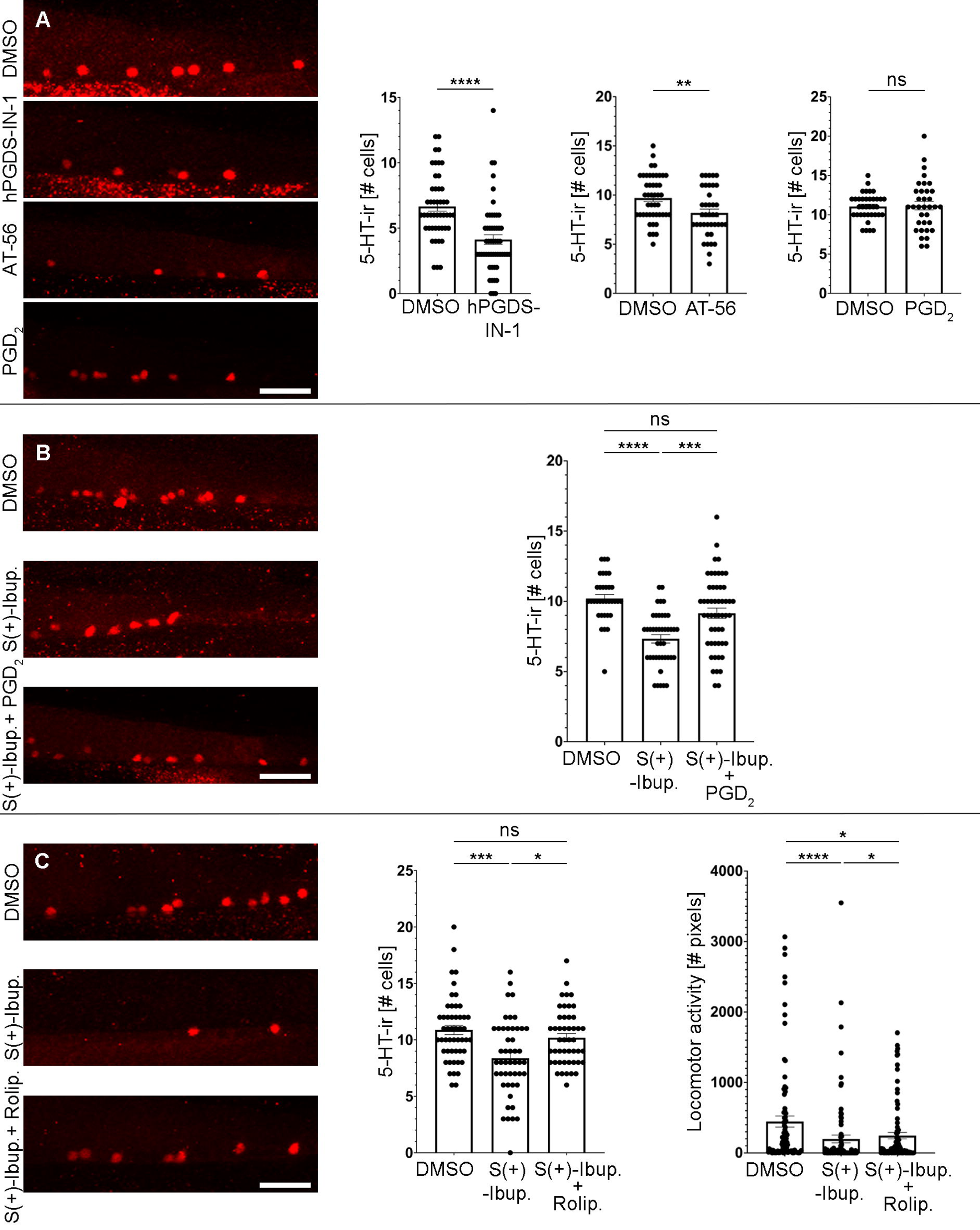
PGDS/PGD_2_/cAMP signalling promotes the generation of serotonergic neurons (red fluorescence) in the ventral spinal cord. A. Treatments with the PGDS inhibitors hPGDS-IN-1 (4.132 ± 0.366 cells, n=53; Mann-Whitney test; p-value <0.0001) and AT-56 (8.179 ± 0.392 cells, n=39; Unpaired t test; p-value = 0.0045) reduced the numbers of serotonergic neurons in the ventral spinal cord as compared to DMSO controls (hPGDS-IN-1 controls: 6.660 ± 0.359 cells, n=47; AT-56 controls: 9.696 ± 0.343 cells, n=46). A treatment with PGD_2_ methyl ester (2.5 µM; 11.18 ± 0.546 cells, n=34; Unpaired t test; p-value = 0.8421) does not significantly change the number of serotonergic neurons in the spinal cord of 4 dpf zebrafish as compared to DMSO controls (11.06 ± 0.282 cells, n=36). B. A co-treatment of S(+)-Ibuprofen (10 µM) with 2.5 µM PGD_2_ methyl ester (9.154 ± 0.364 cells, n=52; Kruskal-Wallis test; p value = 0.071) was able to rescue the inhibitory effects of an S(+)-Ibuprofen treatment (7.333 ± 0.295 cells, n=42; Kruskal Wallis test; p value <0.0001) on the generation of 5-HT-ir spinal cord neurons. Numbers of serotonergic neurons in S(+)-Ibuprofen and PGD_2_ methyl ester treated zebrafish were not significantly different as compared to DMSO controls (10.21 ± 0.283 cells, n=34). C. S(+)-Ibuprofen and Rolipram treated 4 dpf zebrafish showed numbers of 5-HT-ir neurons (10.19 ± 0.361 cells, n=47; Kruskal Wallis test; p value = 0.9081) similar to DMSO controls (10.88 ± 0.411 cells, n=50) and that were significantly higher than S(+)-Ibuprofen treated (8.375 ± 0.489 cells, n=48; Kruskal Wallis test; p value = 0.0006) zebrafish. 4 dpf zebrafish treated with S(+)- Ibuprofen and Rolipram (245.3 ± 45.66 cm, n=89; Kruskal Wallis test; p value = 0.0315) showed increased locomotor activity as compared to S(+)-Ibuprofen (198.7 ± 56.07 cm, n=88; Kruskal Wallis test; p value <0.0001) treated zebrafish (Fig. 3D). However, locomotor activity in S(+)-Ibuprofen and Rolipram treated zebrafish was still significantly lower than in DMSO controls (445.2 ± 79.30 cm, n=85). Rostral is to the right and dorsal to the top in all photomicrographs. Scale bars: 25 µm.

### 3.6 Increasing intracellular cAMP levels rescues the detrimental effects of COX inhibition on spinal cord neurogenesis and locomotion

Unfortunately, the receptor/s for PGD_2_ have not been identified in zebrafish [Zebrafish Information Network (ZFIN) or zebrafish genome searches; see Tsuge et al., 2013], which precludes us from investigating their role in the generation of serotonergic neurons at this point. However, activation of most PG receptors (including PGD_2_ receptors) leads to intracellular changes in cAMP levels (see Nango and Kosuge, 2021). Indeed, most of the PG receptors previously identified in zebrafish (PGI_2_ or PGE_2_ receptors) increase intracellular cAMP levels (Tsuge et al., 2013). Thus, we performed a rescue experiment by treating 2 dpf zebrafish with a combination of S(+)-Ibuprofen and Rolipram (a PDE4 inhibitor that increases intracellular cAMP levels; Lundegaard et al., 2015). S(+)-Ibuprofen and Rolipram treated 4 dpf zebrafish showed numbers of 5-HT-ir neurons similar to DMSO controls and that were significantly higher than S(+)- Ibuprofen treated zebrafish (Fig. 3C). Importantly, 4 dpf zebrafish treated with S(+)- Ibuprofen and Rolipram also showed increased locomotor activity as compared to S(+)- Ibuprofen treated 4 dpf zebrafish (Fig. 3D). However, locomotor activity in S(+)- Ibuprofen and Rolipram treated zebrafish was still significantly lower than in DMSO controls (Fig. 3C). Thus, these rescue experiments indicate that PG signalling might promote neurogenesis in the ventral spinal cord through the intracellular regulation of cAMP levels.

### 3.7 COX2-derived PG signalling promotes mitotic activity in ventral progenitor cells

To investigate the cellular mechanisms by which COX2-derived PGs promote neurogenesis in the developing spinal cord, we analysed levels of apoptotic cell death and mitotic activity in 3 dpf zebrafish after S(+)-Ibuprofen or Meloxicam treatments starting at 2 dpf. We did not find a significant difference in the (very low) number of TUNEL+ (apoptotic) cells in the spinal cord between DMSO controls and S(+)- Ibuprofen or Meloxicam treated 3 dpf zebrafish (Fig. 4A). This indicates: 1) that the treatments with COX inhibitors did not reduce neurogenesis by an increase in apoptotic cell death, and 2) that PGs do not promote neurogenesis by promoting the survival of progenitors or differentiating/differentiated cells. However, S(+)-Ibuprofen or Meloxicam treated 3 dpf zebrafish showed a significant reduction in the number of mitotic (pH3+) cells in the spinal cord (Fig. 4B). Interestingly, when analysing mitotic activity separately for the dorsal and ventral portions of the spinal cord, we observed a significant reduction in the number of mitotic cells only in the ventral region and not in the dorsal region after the S(+)-Ibuprofen or Meloxicam treatments. These results indicate that COX2-derived PGs promote neurogenesis in the ventral spinal cord by promoting mitotic activity in progenitor cells of this region. Future identification of the PG receptor/s implicated in responding to PGD_2_ will allow to determine whether PGD_2_ acts indirectly in an autocrine manner in FP cells or whether it acts in a paracrine manner directly on serotonergic progenitor cells to promote neurogenesis.

**Figure 4.**
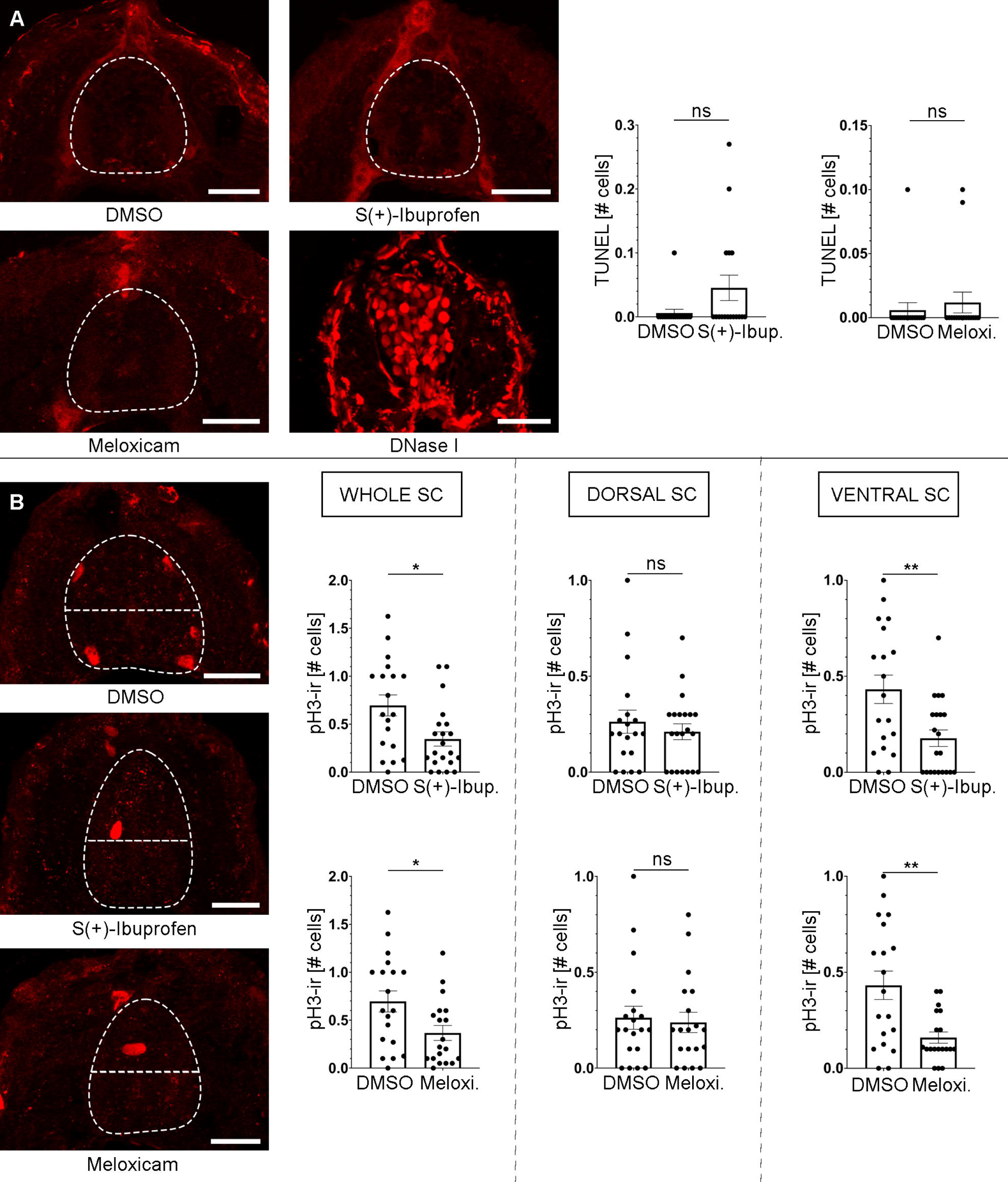
PG signalling promotes mitotic activity in the developing spinal cord of 3 dpf zebrafish. A. Treatments with the COX inhibitors S(+)-Ibuprofen (0.046 ± 0.02 cells, n=17; Mann-Whitney test; p-value = 0.1227) and Meloxicam (0.012 ± 0.008 cells, n=16; Mann-Whitney test; p-value = 0.7341) did not significantly changed the number of TUNEL positive (apoptotic) cells in the spinal cord as compared to DMSO controls (0.006 ± 0.006 cells, n=17). Note the clear TUNEL positive nuclei in a DNase I positive control spinal cord section. B. Treatments with the COX inhibitors S(+)-Ibuprofen (0.345 ± 0.074 cells, n=21; Unpaired t test; p-value = 0.0107) and Meloxicam (0.367 ± 0.078 cells, n=19; Unpaired t test; p-value = 0.0199) significantly changed the number of pH3+ positive (mitotic) cells in the whole spinal cord (SC) as compared to DMSO controls (0.695 ± 0.11 cells, n=19). Treatments with the COX inhibitors S(+)-Ibuprofen (0.210 ± 0.041 cells, n=21; Mann-Whitney test; p-value = 0.9402) and Meloxicam (0.238 ± 0.053 cells, n=29; Mann-Whitney test; p-value = 0.8211) did not significantly changed the number of pH3+ positive cells in the dorsal spinal cord as compared to DMSO controls (0.263 ± 0.060 cells, n=19). Treatments with the COX inhibitors S(+)- Ibuprofen (0.177 ± 0.043 cells, n=21; Unpaired t test; p-value = 0.0042) and Meloxicam (0.16 ± 0.029 cells, n=29; Unpaired t test; p-value = 0.0016) significantly changed the number of pH3+ positive cells in the ventral spinal cord as compared to DMSO controls (0.432 ± 0.074 cells, n=19). Dorsal is to the top in all photomicrographs of the larval transverse sections. The dashed lines indicate the border of the spinal cord in A and B and, also the separation between the ventral and dorsal portions of the spinal cord in B. Scale bars: 20 µm.

As indicated above, there is very limited data on the role of PGs in neurogenesis and the available data comes only from studies in vitro or in adult rodents. A previous study showed that in adult rats a treatment with Sulprusone (a PGE_2_ analogue) increased cell proliferation in the dentate gyrus of the hippocampus (Uchida et al., 2002). Interestingly, in the adult mouse hippocampus, Rolipram administration also increases cell proliferation and the generation of mature granule cells (Nakagawa et al., 2002). Our Rolipram experiments (see above) also suggest that PG signalling promotes mitotic activity in spinal cord progenitor cells by increasing intracellular cAMP levels. Noteworthily, intracellular cAMP regulates the response to Shh signalling through protein kinase A (see Saad and Hipfner, 2021). In vitro work has also shown that PGE_2_ interacts with Wnt signalling through PKA to promote cell proliferation in neuroectodermal (NE-4C) stem cells (Wong et al., 2014). Moreover, the expression of Wnt signalling genes is altered in COX2 knockout mice (Rai-Bhogal et al., 2018a) and in the offspring after maternal exposure to PGE_2_ (Rai-Bhogal et al., 2018b). Thus, a possible interaction between PGD_2_ and Shh or Wnt signalling to regulate the behaviour of progenitor cells in the spinal cord is suggested and deserves further investigation.

Although our study reveals that COX2 FP-derived PGD_2_ promotes neurogenesis in the ventral spinal cord by promoting mitotic activity in progenitor cells, we should not discard a possible role for PG signalling in neuronal differentiation. In vitro studies have shown that PGE_2_ promotes the differentiation of NE-4C cells into neuronal-linage cells (Wong et al., 2016). Also, in vitro work using the mouse neuroblastoma and spinal cord motor neuron fusion cell line (NSC-34), has shown that PGE_2_ increases the percentage of neurite-bearing cells (Nango et al., 2017) and the differentiation of NSC-34 cells into functional motor neurons (Nango et al., 2020a). In the same cell line, PGD_2_ also increased the proportion of cells with neurites and neurite length in NSC-34 cell cultures (Nango et al. 2020b). Therefore, a possible role for PGD_2_ signalling in promoting neuronal differentiation in the developing spinal cord should not be excluded. A recent review article stated that a main limitation on the study of the roles of PGs in neurogenesis was the lack of in vivo studies (Nango and Kosuge, 2021). Our in vivo work in zebrafish starts to fill this gap in our knowledge by showing that PGD_2_ signalling promotes neurogenesis in the developing spinal cord.

## 4. Conclusions

Our data shows that COX2 and PGDS/PGD_2_ activity in the FP promotes neurogenesis in the ventral spinal cord by promoting mitotic activity in progenitor cells. Future work should attempt to study the possible interactions between the regulation of intracellular cAMP levels by PGs and Shh or Wnt signalling during spinal cord neurogenesis. The lack of identification of the PGD_2_ receptor/s in zebrafish precluded us from determining whether PGD_2_ acts in an autocrine manner on FP cells (indirect regulation of neurogenesis) or in a paracrine manner (direct regulation of serotonergic neuronal progenitors). Thus, finding the receptor/s mediating PGD_2_ signalling in zebrafish would be of great interest. In any case, our research uncovers a previously unknown regulatory pathway for neurogenesis in the spinal cord of vertebrates, which may also have implications for the clinical use of NSAIDs. If PG signalling has similar effects on early neurogenesis in mammals (including humans) our results could have far-reaching consequences for the use of NSAIDs in pregnant women/infants. Moreover, there are ongoing studies and clinical trials testing the use of Ibuprofen as a Rho inhibitor to promote recovery after spinal cord injury (Watzlawick et al., 2014; Kopp et al., 2012, 2016). Thus, our results should be considered due to possible effects of Ibuprofen or other NSAIDs on regenerative neurogenesis or in cell replacement therapies that are also being tested in the context of spinal cord injury (see Ribeiro et al., 2023).

## Author contributions

**Laura González-Llera:** Investigation, Formal analysis, Visualization, Writing- Reviewing and Editing. **Daniel Sobrido-Cameán:** Investigation, Formal analysis, Writing- Reviewing and Editing. **Ana Quelle-Regaldie:** Investigation, Formal analysis, Writing- Reviewing and Editing. **Laura Sánchez:** Funding acquisition, Resources, Writing- Reviewing and Editing. **Antón Barreiro-Iglesias:** Conceptualization, Methodology, Project administration, Funding acquisition, Supervision, Formal analysis, Writing- Original draft preparation.

## Data availability statement

Raw data and materials not included within the article are available from the authors upon reasonable request.

## 5. Acknowledgements

We would like to thank the Servizo de Microscopía of the University of Santiago de Compostela and Dr. Mercedes Rivas Cascallar for confocal microscope facilities and technical help.

## Supplemental information

**Supplementary File 1.** Results from the primary drug screen.

**Supplementary File 2.** Results from the secondary drug screen. Drugs mentioned in the manuscript are indicated in red.

**Supplementary File 3.** Results from the treatments with COX2 inhibitors of the LOPAC library.

**Supplementary Figure 1.** A. Schematic drawing showing the experimental design and morpholino and drug treatments’ time windows. B. Schematic of the drug screening protocol. Abbreviations: IF, immunofluorescence; MOs, morpholinos; SC, spinal cord.

**Supplementary Figure 2.** Graphs from the scRNAseq atlas (each individual cell is indicated by a light blue dot) of developing zebrafish (Farnsworth et al., 2020) showing the expression of *ptgs2a* (A), *ptgdsb.2* (B) and *ptgdsb.1* (C) in FP cells (cluster 176; cells of this cluster are indicated by black circles). FP cells expressing these genes are color coded according to the legends on the right.

