## Supplementary figures and images for "An in vivo drug screen reveals that cyclooxygenase 2-derived prostaglandin D_2_ promotes spinal cord neurogenesis"

### Supplementary figure 1

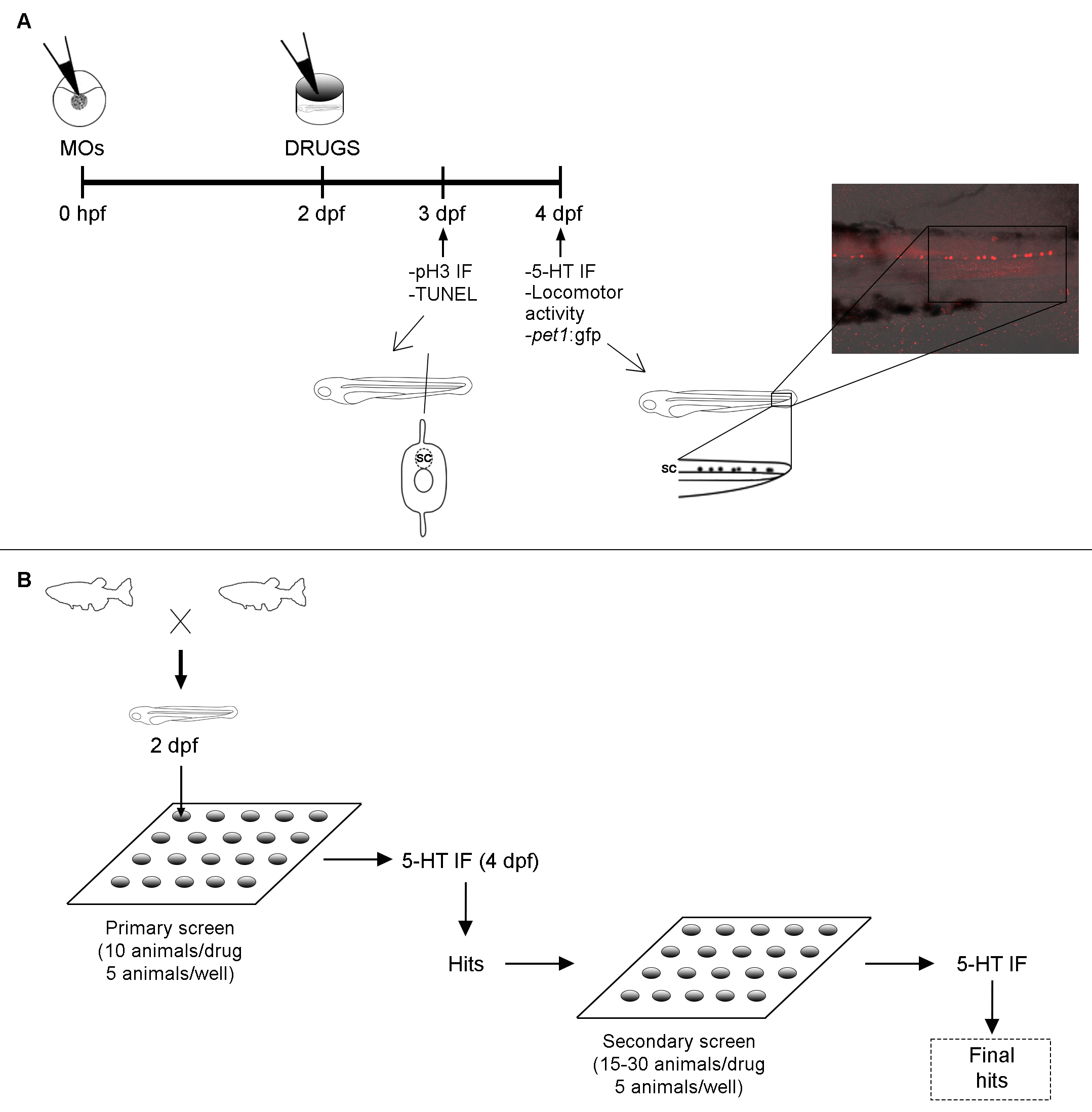

### Supplementary figure 2

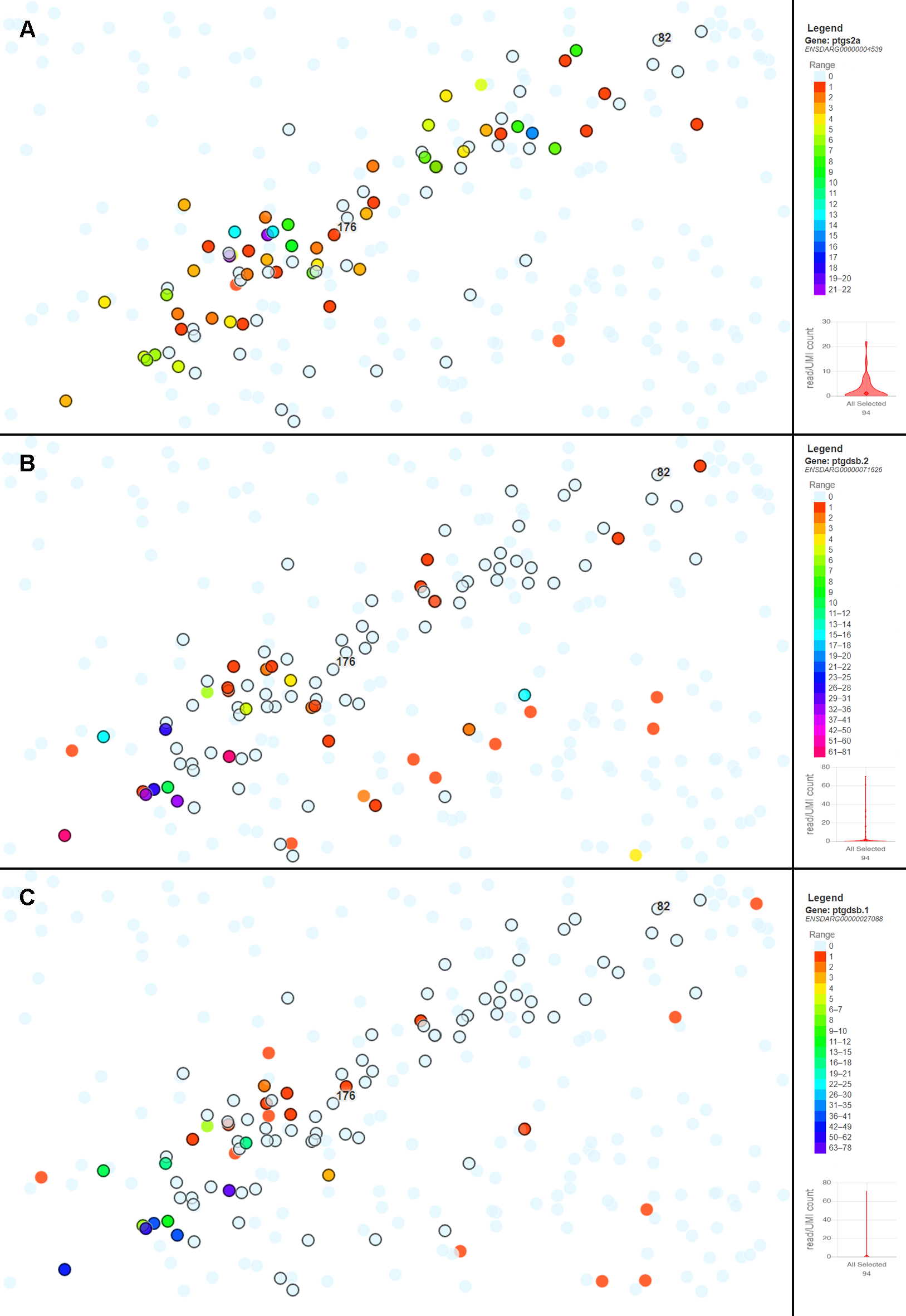
